# High abundance of transcription regulators compacts the nucleoid in *Escherichia coli*

**DOI:** 10.1101/2022.01.19.477023

**Authors:** Cihan Yilmaz, Karin Schnetz

## Abstract

In enteric bacteria organization of the circular chromosomal DNA into a highly dynamic and toroidal shaped nucleoid involves various factors such as DNA supercoiling, nucleoid-associated proteins (NAPs), the structural maintenance of chromatin (SMC) complex, and macro-domain organizing proteins. Here we show that ectopic expression of transcription regulators at high levels leads to nucleoid compaction. This serendipitous result was obtained by fluorescence microscopy upon ectopic expression of the transcription regulator and phosphodiesterase PdeL of *Escherichia coli* of a strain expressing the mCherry-tagged HU-α subunit (HupA) for nucleoid staining. Nucleoid compaction by PdeL depends on DNA-binding, but not on its enzymatic phosphodiesterase activity. Nucleoid compaction was also observed upon high-level ectopic expression of the transcription regulators LacI, RutR, RcsB, LeuO and Cra, which range from single target gene regulators to global regulators. In case of LacI its high-level expression in presence of the gratuitous inducer IPTG also led to nucleoid compaction indicating that compaction is caused by unspecific DNA-binding. In all cases nucleoid compaction correlated with misplacement of the FtsZ ring and loss of MukB foci, a subunit of the SMC complex. Thus, high levels of several transcription regulators cause nucleoid compaction with consequences on transcription, replication, and cell division.

**Importance:** The bacterial nucleoid is a highly organized and dynamic structure for simultaneous, transcription, replication and segregation of the bacterial genome. Compaction of the nucleoid and disturbance of DNA segregation and cell division by artificially high levels of transcription regulators, as described here, reveals that an excess of DNA-binding protein disturbs nucleoid structuring. The results suggest that ectopic expression levels of DNA-binding proteins for genetic studies of their function but also for their purification should be carefully controlled and adjusted.

## Introduction

Regulation of transcription in *Escherichia coli* involves a repertoire of approximately 300 transcription regulators of which more than 90% have been functionally validated (1–3). These transcription regulators include single target regulators such as LacI solely regulating the *lac* operon, local regulators with up to 50 target genes as for example RutR, a pyrimidine utilization repressor (4), and global regulators such as the catabolite activator repressor protein Cra and the pleiotropic regulator LeuO with more than 100 targets (5, 6). Nucleoid-associated proteins (NAPs) constitute a further group of DNA-binding proteins; they are abundant, relevant for organization of the genomic DNA as a nucleoid, and participate in the regulation of hundreds of targets genes (7).

PdeL, carrying a N-terminal FixJ/NarL/LuxR-type DNA-binding domain and a C-terminal EAL-type c-di-GMP-specific phosphodiesterase domain, is one of the validated transcriptional regulators with a small number of target loci including the *fliFGHIK* operon, *sslE*, and *pdeL* itself (8, 9). However, the physiological function of PdeL remains an open question, as a *pdeL* deletion mutant has no significant phenotype at least at the laboratory growth conditions (9). At these growth conditions expression of the *pdeL* gene and concomitantly PdeL protein levels are low, due to repression of *pdeL* by the abundant nucleoid-associated and global repressor protein H-NS (9, 10). However, moderately elevated PdeL levels using plasmids or up-regulated *pdeL* mutants disclosed its function as transcription regulator and as active c-di-GMP specific phosphodiesterase (8, 9). Furthermore, PdeL, as a dual function protein may represent a trigger enzyme whose role as transcription regulator is controlled by c-di-GMP via the phosphodiesterase domain (11).

Considering the dual functions of PdeL we used a fluorescent protein fusion, PdeL-mVenus, provided by a pBAD-derived plasmid to analyze its cellular localization. Serendipitously, we found that ectopic expression of PdeL causes nucleoid compaction even upon weak induction of the *P*_*ara*_ promoter directing expression of *pdeL-mVenus*. Weak induction nonetheless led to high levels of PdeL, which is a very stable protein. Further, other transcription regulators (LacI, RutR, RcsB, LeuO, and Cra) all cause compaction of the nucleoid as well, when expressed and synthesized at similarly high levels. The data indicate that the mere occupation of the genomic DNA by an abundant DNA-binding protein can have severe effects on nucleoid structuring.

## Results

### PdeL is nucleoid-associated and causes nucleoid compaction

Here we addressed whether the dual function protein PdeL is predominantly nucleoid-associated or localized otherwise. To this end, PdeL-mVenus fusions were provided by plasmids under control of the *P*_*ara*_ promoter, and the cellular localization was analyzed by fluorescence microscopy in *E. coli hupA-mCherry* Δ*pdeL* strain U159 (Fig. 1A, Tables 1 and 2). In this strain gradual induction of *P*_*ara*_ by arabinose is applicable due to modification of the arabinose regulon, as described (12, 13), while HupA-fluorescent protein fusions are well-established markers for nucleoid imaging (14, 15). Cellular localization of PdeL-mVenus and of PdeL_HTH5M_*-*mVenus, a mutant defective in DNA-binding as a control, was analyzed (9). Note that chromosomal encoded PdeL-mVenus is not detectable by fluorescence microscopy due to low expression of the *pdeL* gene (9).

**Table 1:**
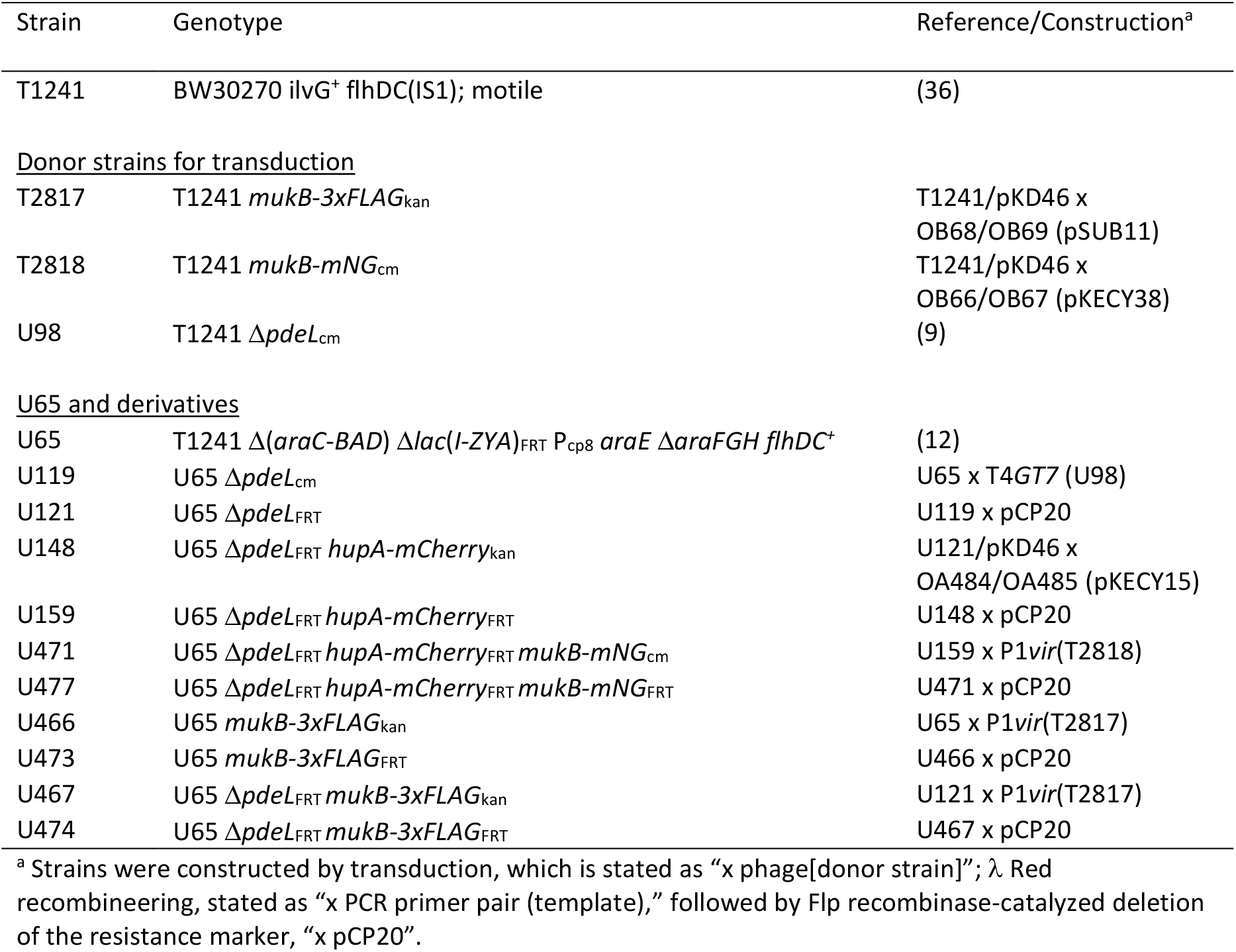
E. coli K12 strains

**Table 2.**
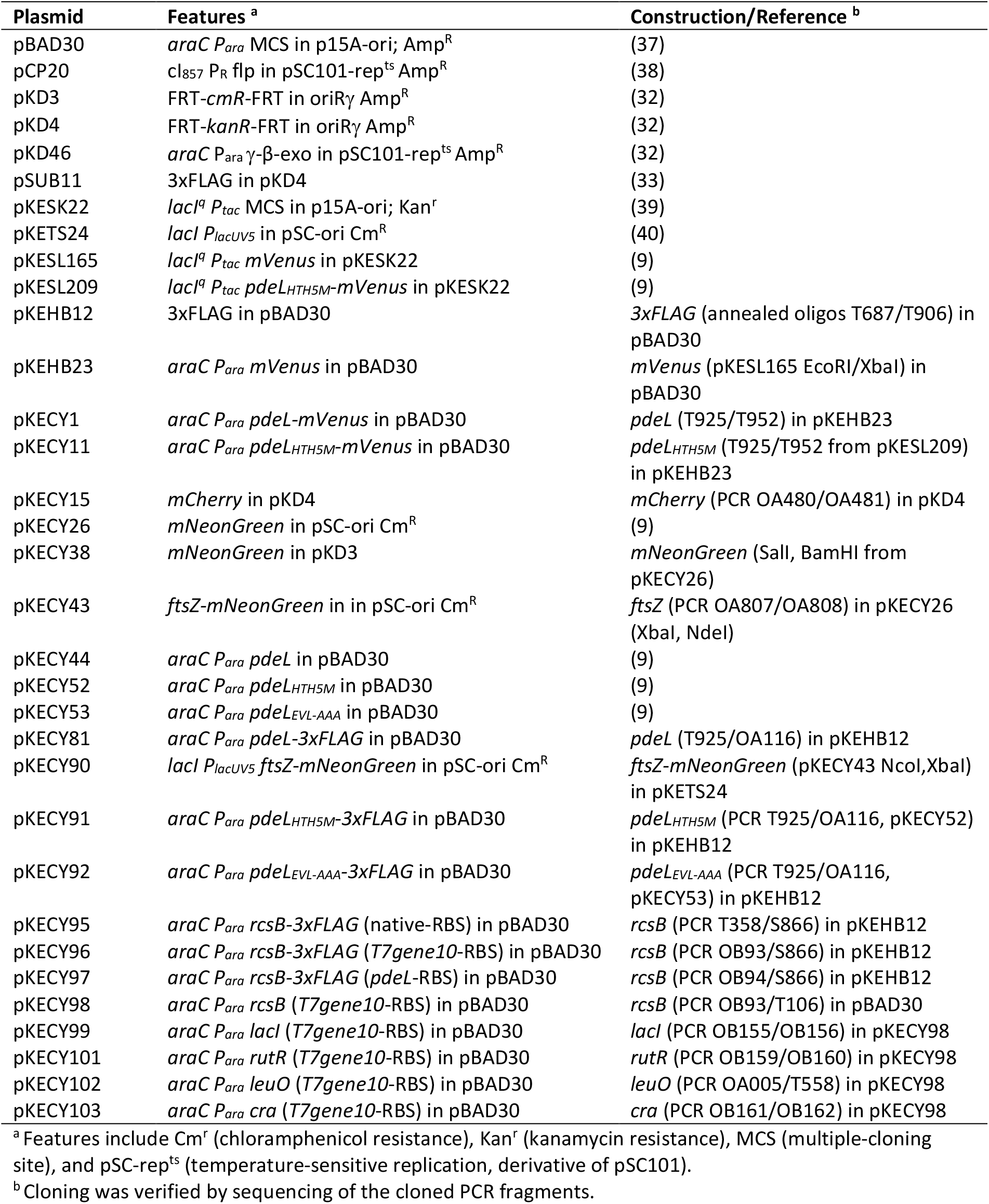
Plasmids.

**Figure 1.**
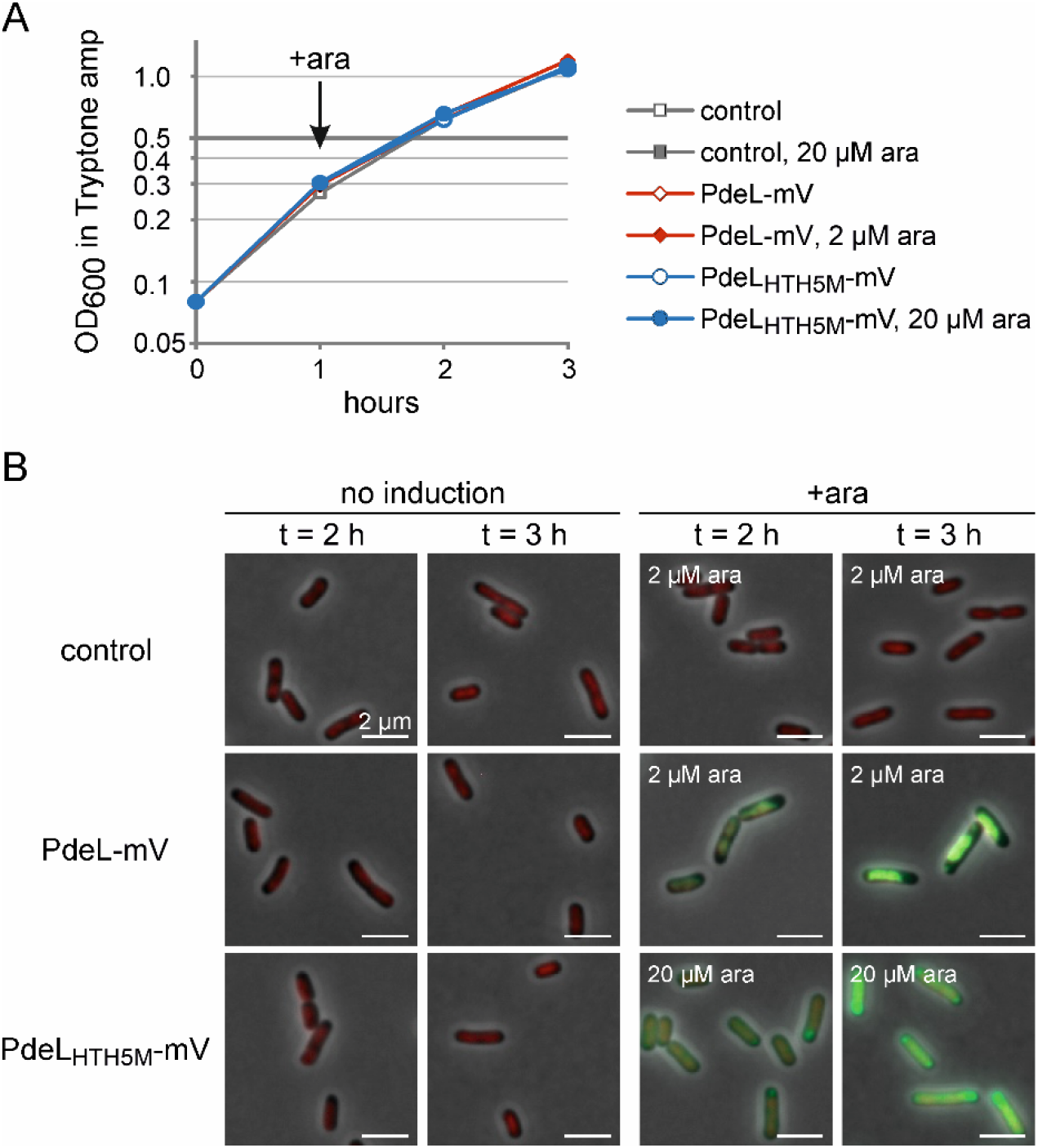
PdeL is nucleoid-associated. (A) Growth of transformants of *hupA-mCherry* Δ*pdeL* strain U159 with plasmids pBAD30 (control), pKECY1 (PdeL-mV) and pKECY11 (PdeL_HTH5M_*-*mV), respectively. Cultures were inoculated to OD_600_ 0.08 and grown in tryptone ampicillin medium. After one hour of growth (t = 1 h), *P*_*ara*_ directed expression was induced with L-arabinose using a final concentration of 2 µM in case of *pdeL-mVenus* and 20 µM in case of *pdeL*_*HTH5M*_*-mVenus*. With these inducer concentrations steady state protein levels are similar as tested using 3xFLAG variants of PdeL and PdeL_HTH5M_ (Fig. S1). (B) Composite fluorescent microscopy images of representative bacteria with fluorescence of HupA-mCherry shown in red and of PdeL-mVenus or PdeL_HTH5M_-mVenus (in green). Samples were harvested from un-induced and induced cultures at t = 2 h and t = 3 h. The scale bar corresponds to 2 µm. Full images are shown in supplementary Figure S2.

First, we determined the L-arabinose concentration needed for inducing the synthesis of equal amounts of PdeL and the DNA-binding defective control PdeL_HTH5M_ by using 3xFLAG alleles carried on the same plasmidic expression system as used for fluorescence microscopy (Fig. S1). PdeL-3xFLAG gene expression was induced with 2µM L-arabinose, while expression of PdeL_HTH5M_-3xFLAG was induced by addition of increasing concentrations of L-arabinose. The amount of PdeL-3xFLAG (induced by addition of 2 µM L-arabinose) and PdeL_HTH5M_-3xFLAG (induced with 20 µM L-arabinose) were similar (Fig. S1). The low concentration of L-arabinose that is required for synthesis of significant amounts of PdeL indicates that PdeL synthesis is efficient and that the PdeL protein is stable. Determination of the protein stability of PdeL and PdeL_HTH5M_ showed that PdeL-3xFLAG was stable for 160 minutes after inhibition of translation by 100 µg/ml chloramphenicol (Fig. S1). In contrast, the PdeL_HTH5M_-3xFLAG steady state level was lower and its level decreased approximately 2-fold in the 160 minutes after inhibition of translation, which suggests that PdeL_HTH5M_ is less stable than PdeL (Fig. S1). Note that protein stability was determined without induction to keep PdeL-levels sufficiently low for quantification (Fig. S1). The result of the protein stability assay is in accordance with the different concentrations of L-arabinose that are required for similar steady state protein levels of PdeL-3xFLAG and PdeL_HTH5M_-3xFLAG.

For fluorescence microscopy plasmidic *P*_*ara*_*-*directed expression of *pdeL-Venus* and *pdeL*_*HTH5M*_*-mVenus* was induced by L-arabinose after one hour of growth (t = 1h), and samples were harvested after one and two additional hours of growth (at t=2h and t=3h) later (Fig. 1). PdeL-mVenus fluorescence was apparent after one hour of induction and localized to the whole nucleoid (Fig. 1B and Fig. S2). In addition, nucleoid-association was accompanied by nucleoid compaction, formation of nucleoid free areas near the cell poles, and an enlargement of the bacteria. In contrast, PdeL_HTH5M_-mVenus was located diffusely in the cell, and after two hours of induction, aggregates of PdeL_HTH5M_-mVenus near the cell pole became apparent. Taken together, the data suggest that the transcription regulator PdeL is nucleoid-associated and they indicate that DNA-binding by high PdeL levels cause nucleoid compaction.

### PdeL and RcsB compact the nucleoid and affect localization of MukB-mNeonGreen

PdeL-mVenus leads to nucleoid compaction close to midcell. Since the nucleoid structure is changed, we tested localization of MukB, a subunit of the structural maintenance of chromosome (SMC) complex and a marker for *oriC* localization, using chromosomally encoded MukB-mNeonGreen (MukB-mNG) (16–19). In addition, FtsZ-mNG was used as a marker for septum formation (for results see below). In this experimental approach, transformants expressing non-tagged PdeL, the DNA-binding defective, PdeL_HTH5M_, and an enzymatically inactive PdeL_EVL-AAA_ were studied (Fig. 2A). PdeL_EVL-AAA_ is enzymatically inactive due to mutation of the conserved c-di-GMP-specific EVL motif to three alanine residues. In addition to PdeL, high level expression of the two-component response regulator RcsB was included, which like PdeL carries a FixJ/NarL/LuxR-type DNA-binding domain, to analyze whether nucleoid compaction by high protein levels is PdeL-specific.

**Figure 2.**
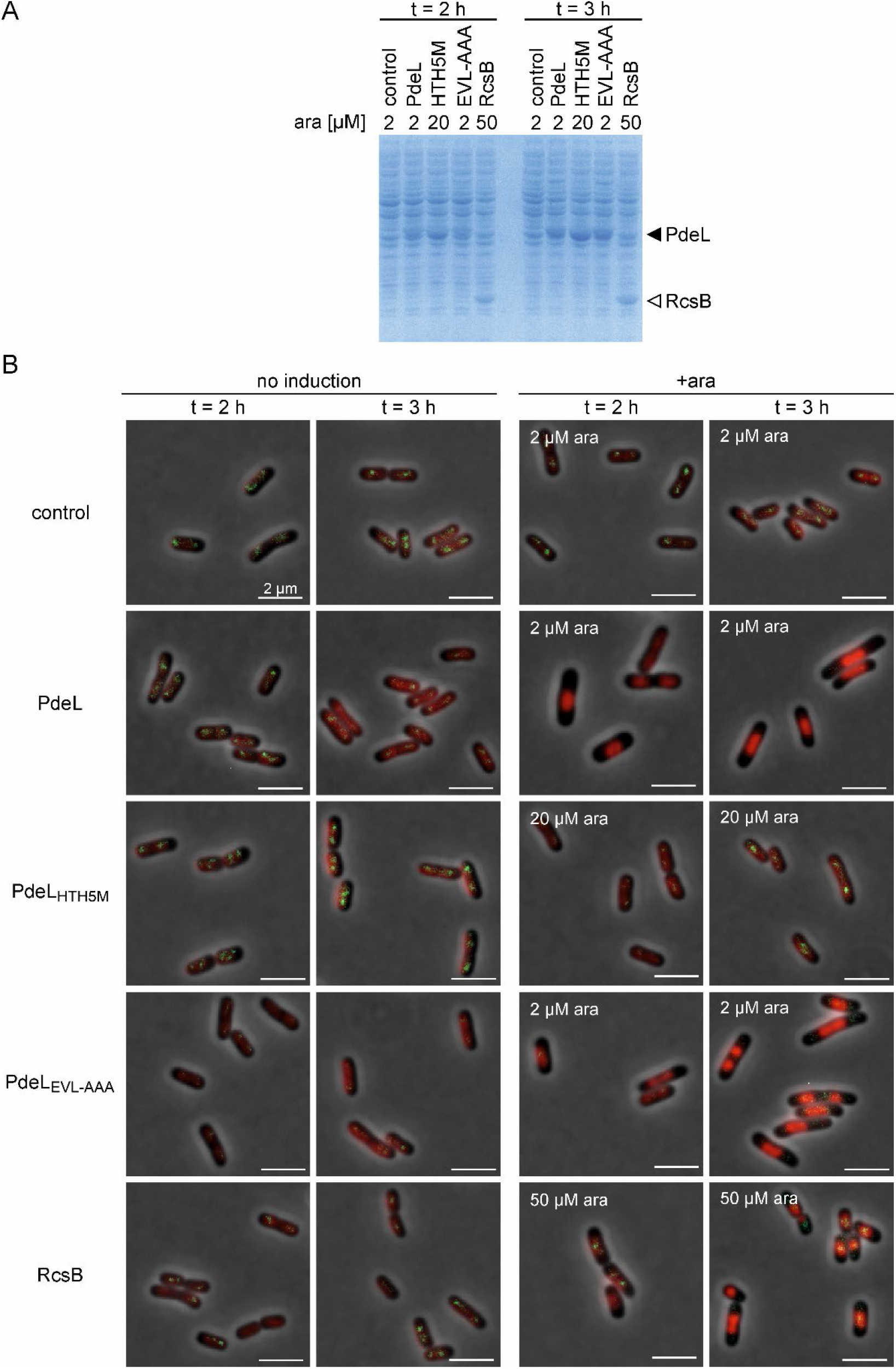
High levels of PdeL, PdeL_EVL-AAA_ and RcsB cause nucleoid compaction and loss of MukB-mNG foci. Transformants of *hupA-mCherry mukB-mNG* Δ*pdeL* strain U477 with plasmids pBAD30 (control), pKECY44 (PdeL), pKECY52 (PdeL_HTH5M_), pKECY53 (PdeL_EVL-AAA_), and pKECY98 (RcsB), respectively, were grown in tryptone ampicillin medium. After one hour of growth *P*_*ara*_ directed expression of *pdeL* and its mutants as well as of *rcsB* was induced by addition of L-arabinose, as indicated. Samples were harvested after 2 and 3 hours of growth (t = 2h, t= 3h). (A) Protein levels were analyzed by 15% SDS-PAGE and Coomassie staining. Bands corresponding to PdeL proteins and to RcsB are indicated, by filled and open triangles, respectively. (B) Representative sections of microscopy images of transformants expressing no protein (control), PdeL, PdeL_HTH5M_, PdeL_EVL-AAA_, and RcsB, respectively. HupA-mCherry fluorescence is shown in red and MukB-mNeonGreen foci are shown in green. Full size microscopy images are shown in Fig. S4.

First, synthesis of RcsB levels that are similar to the levels of PdeL were established using plasmids carrying 3xFLAG-tagged *rcsB* alleles under *P*_*ara*_ control (Fig. S3). In these plasmids the efficiency of translation of *rcsB-3xFLAG* was varied. RcsB-3xFLAG levels were lowest with its native ribosomal binding site (RBS), higher when *rcsB* was fused to *pdeL’s* RBS, and similarly high as PdeL levels when using the phage *T7 gene10* RBS and 50 µM of L-arabinose for induction (Fig. S3).

Next, fluorescence microscopy was performed using transformants of *hupA-mCherry mukB-mNG* Δ*pdeL* strain U477 with plasmids coding for PdeL, PdeL_HTH5M_, PdeL_EVL-AAA_, and RcsB, respectively. These transformants synthesized approximately equal amounts of PdeL, PdeL_HTH5M_, PdeL_EVL-AAA_ and RcsB upon induction by L-arabinose, as shown by SDS-PAGE (Fig. 2A). Induction of high levels of PdeL, PdeL_EVL-AAA_, and RcsB synthesis caused nucleoid compaction to midcell and a moderate increase of the cell length (Fig. 2B). Furthermore, one or two MukB-mNG foci close to *oriC* were detectable without induction and in the control (Fig. 2B and Fig. S4), as previously shown for fluorescent protein MukB fusions (17). MukB-mNG foci disappeared upon induction of PdeL, PdeL_EVL-AAA_, and RcsB expression, but were not changed in case of the DNA-binding deficient PdeL_HTH5M_ (Fig. 2B). The disappearance of MukB-mNG foci was not caused by a change in the MukB protein levels, which remained constant (Fig. S5). Taken together, high protein levels of PdeL and RcsB, but not of the DNA-binding defective PdeL_HTH5M_ mutant led to nucleoid compaction and loss of MukB-mNG foci.

### High levels of PdeL and RcsB proteins lead to misplacement of the FtsZ ring

As a second marker we tested whether localization of cell division protein FtsZ is affected by synthesis of high levels of PdeL and RcsB. For visualization of the Z-ring in *ftsZ*^*+*^ background, FtsZ-mNG was provided by a low-copy plasmid carrying a *P*_*lacUV5*_ *ftsZ-mNG* cassette, as described (20). The low activity of the non-induced *P*_*lacUV5*_ promoter was sufficient for production of detectable amounts of FtsZ-mNG. Fluorescence microscopy was performed with double transformants of *hupA-mCherry* Δ*pdeL* strain U159 with the *ftsZ-mNG* plasmid and compatible *pdeL* or *rcsB* carrying plasmids (Fig. 3). Depending on the progression of the cell cycle, FtsZ-mNG was detectable at midcell forming the Z-ring, in case of the control and the PdeL_HTH5M_ DNA-binding mutant (Fig. 3 and Fig. S6). Localization of FtsZ-mNG was different when PdeL and when RcsB were expressed at high levels. In these cases, the FtsZ was displaced from mid-cell towards the quarter positions which are devoid of the nucleoid (Fig. 3 and Fig. S6). Taken together, the FtsZ ring is misplaced by high levels of PdeL, PdeL_EVL-AAA_, and RcsB, but not by the DNA-binding defective PdeL_HTH5M_ (Fig. 3 and Fig. S6).

**Figure 3:**
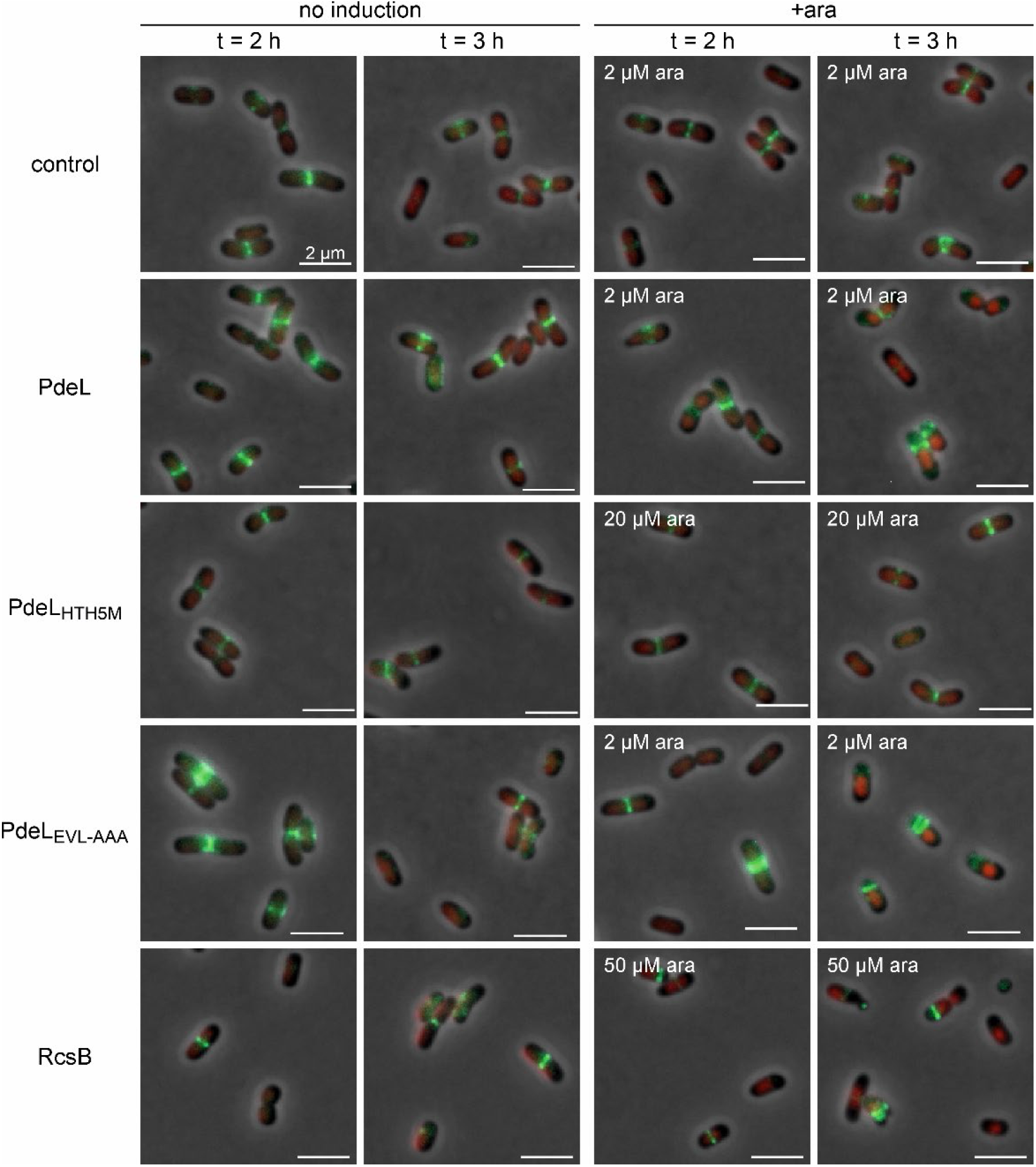
PdeL, PdeL_EVL-AAA_ and RcsB misplace FtsZ rings. For visualization of the Z-ring a C-terminally mNeonGreen tagged FtsZ variant, FtsZ-mNG, was ectopically expressed under the control of *P*_*lacUV5*_ promoter using low copy plasmid pKECY90. Co-transformants of *hupA-mCherry* Δ*pdeL* strain U159 with plasmid pKECY90 (*P*_*lacUV5*_ *ftsZ-mNG*) and plasmids pBAD30 (control), pKECY44 (PdeL), pKECY52 (PdeL_HTH5M_), pKECY53 (PdeL_EVL-AAA_), and pKECY98 (RcsB), respectively, were grown in tryptone ampicillin chloramphenicol medium. After one hour of growth *P*_*ara*_ directed expression was induced by L-arabinose, as indicated. Samples were harvested at t = 2 h and t = 3 h of growth. Shown are representative sections of composite microscopy images of transformants with HupA-mCherry (in red) and FtsZ-mNG (in green). Contrast settings for RcsB images were adjusted for better visibility of FtsZ-mNG to 180-700 (mNG) and 200-2000 (mCherry). Full size images are shown in Fig. S6.

### High levels of the transcription regulators LacI, RutR, LeuO, and Cra also cause nucleoid compaction

Since high levels of both PdeL and RcsB affect the nucleoid structure and localization of MukB and FtsZ, we tested additional DNA-binding proteins. This included the single target transcription regulator LacI, the local regulator RutR, and the global regulators LeuO and Cra (6). Plasmidic *P*_*ara*_ directed expression of these transcription regulators was adjusted to obtain approximately equal amounts of each protein, as validated by SDS-PAGE (Fig. S7). Fluorescence microscopy demonstrated that high levels of all transcription regulators, LacI, RutR, LeuO, and Cra, caused nucleoid compaction as well (Fig. 4 and Fig. S8). In addition, we also analyzed whether LacI in presence of IPTG causes the same phenotype. Specific DNA-binding of LacI is inhibited by IPTG (21). Non-specific DNA-binding of high levels of LacI-IPTG was sufficient to cause compaction of the nucleoid (Fig. 5 and S9). In all cases the number of MukB-mNG foci was significantly lower (Figs. 4, 5, S8, and S10). Taken together, all tested transcription regulators LacI, RutR, LeuO, and Cra caused nucleoid compaction and a decrease of MukB-mNG foci similar to PdeL and RcsB.

**Figure 4:**
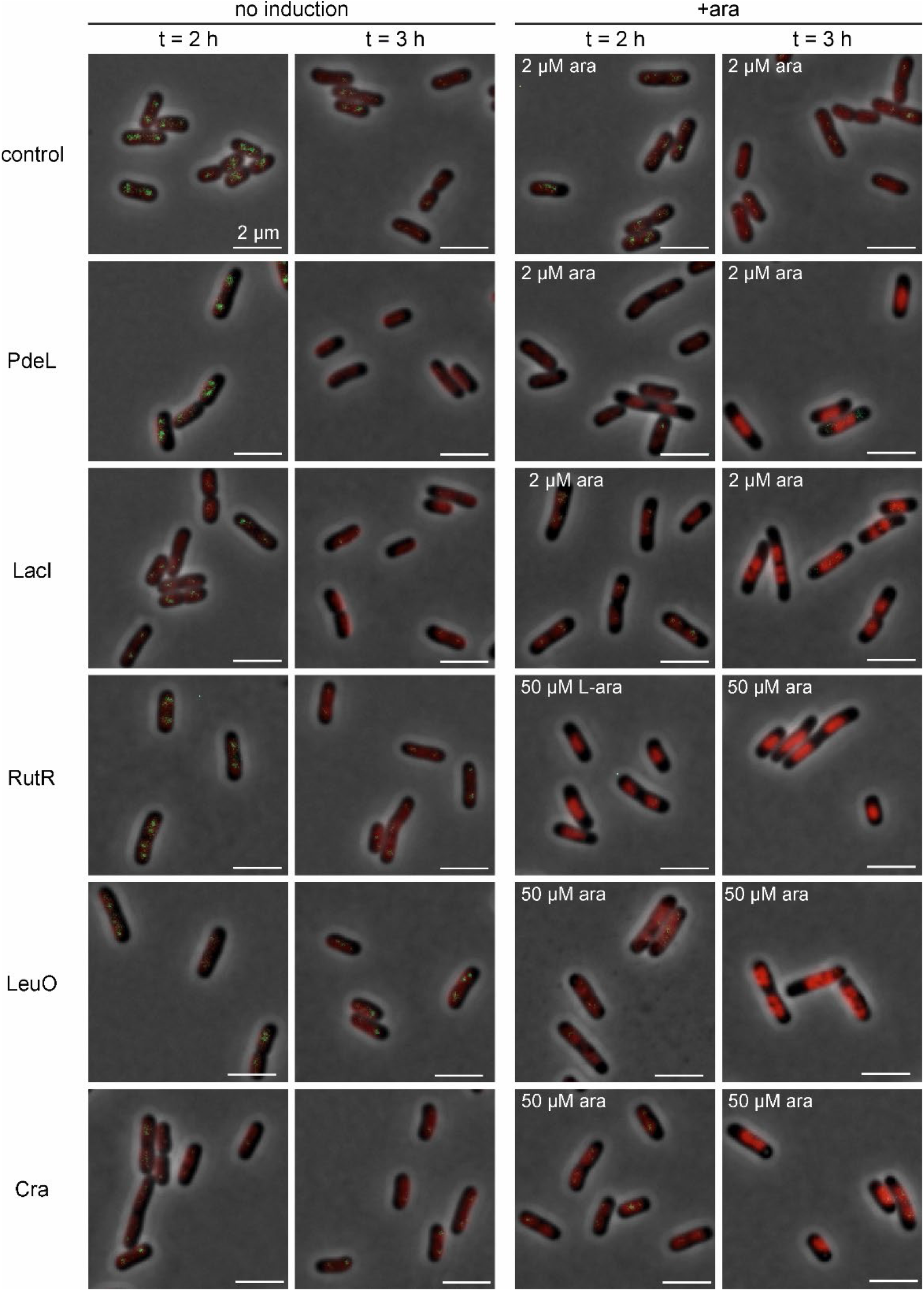
High levels of the transcription regulators LacI, RutR, LeuO, and Cra cause nucleoid compaction and loss of MukB foci. Transformants of *hupA-mCherry mukB-mNG* Δ*pdeL* strain U477 with plasmids pBAD30 (control), pKECY44 (PdeL), pKECY99 (LacI), pKECY101 (RutR), pKECY102 (LeuO), and pKECY103 (Cra) were grown in tryptone ampicillin medium. After one hour of growth, *P*_*ara*_ directed expression of transcription regulators was induced with L-arabinose, as indicated. Expression of similar amounts of the transcription regulators was analyzed SDS-PAGE (Fig. S7). Shown are representative composite microscopy images (full size images are shown in Fig. S8).

**Figure 5:**
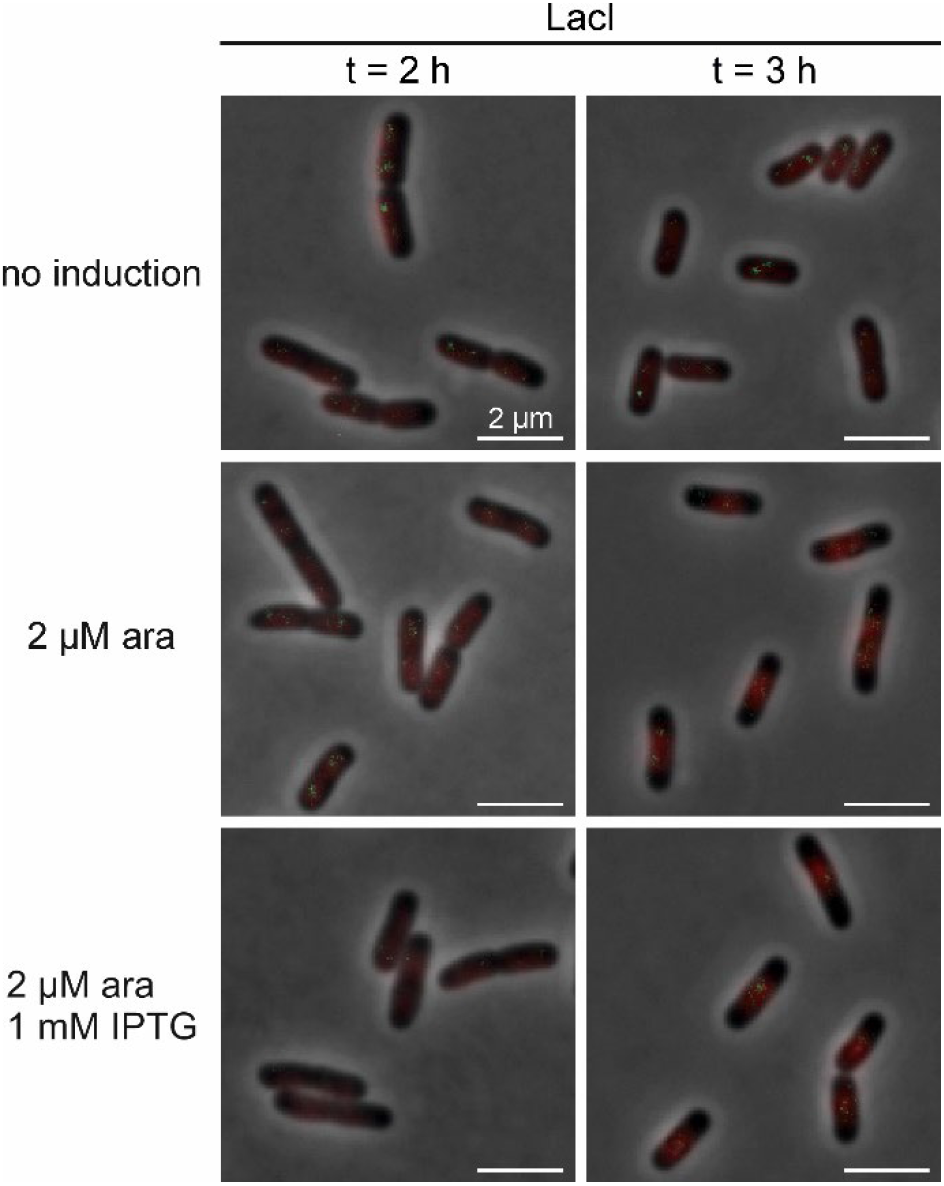
Non-specific DNA-binding by LacI causes nucleoid compaction. To test whether non-specific DNA-binding by LacI causes nucleoid compaction IPTG was added as well to cultures of transformants of *hupA-mCherry mukB-mNG* Δ*pdeL* strain U477 with plasmid pKECY99 (LacI). After one hour of growth in tryptone ampicillin medium *lacI* expression was induced with 2 µM L-arabinose and IPTG was supplemented to a final concentration of 1 mM, where indicated. Samples were harvested after one (t = 2h) and two (t = 3h) hours of growth. Shown are representative composite microcopy images; full size images are shown in Fig. S9.

For RutR, LeuO and Cra we also tested FtsZ localization using low-copy plasmid carrying *ftsZ-mNG* under control of the LacI-regulated *lacUV5* promoter (Fig. 6 and S10). In this experiment LacI could not be included, since high levels of LacI led to complete inhibition of *ftsZ-mNG* expression. Fluorescence microscopy of the transformants expressing high levels of RutR, LeuO and Cra, respectively, caused misplacement of the FtsZ similarly as PdeL and RcsB (Fig. 6).

**Figure 6:**
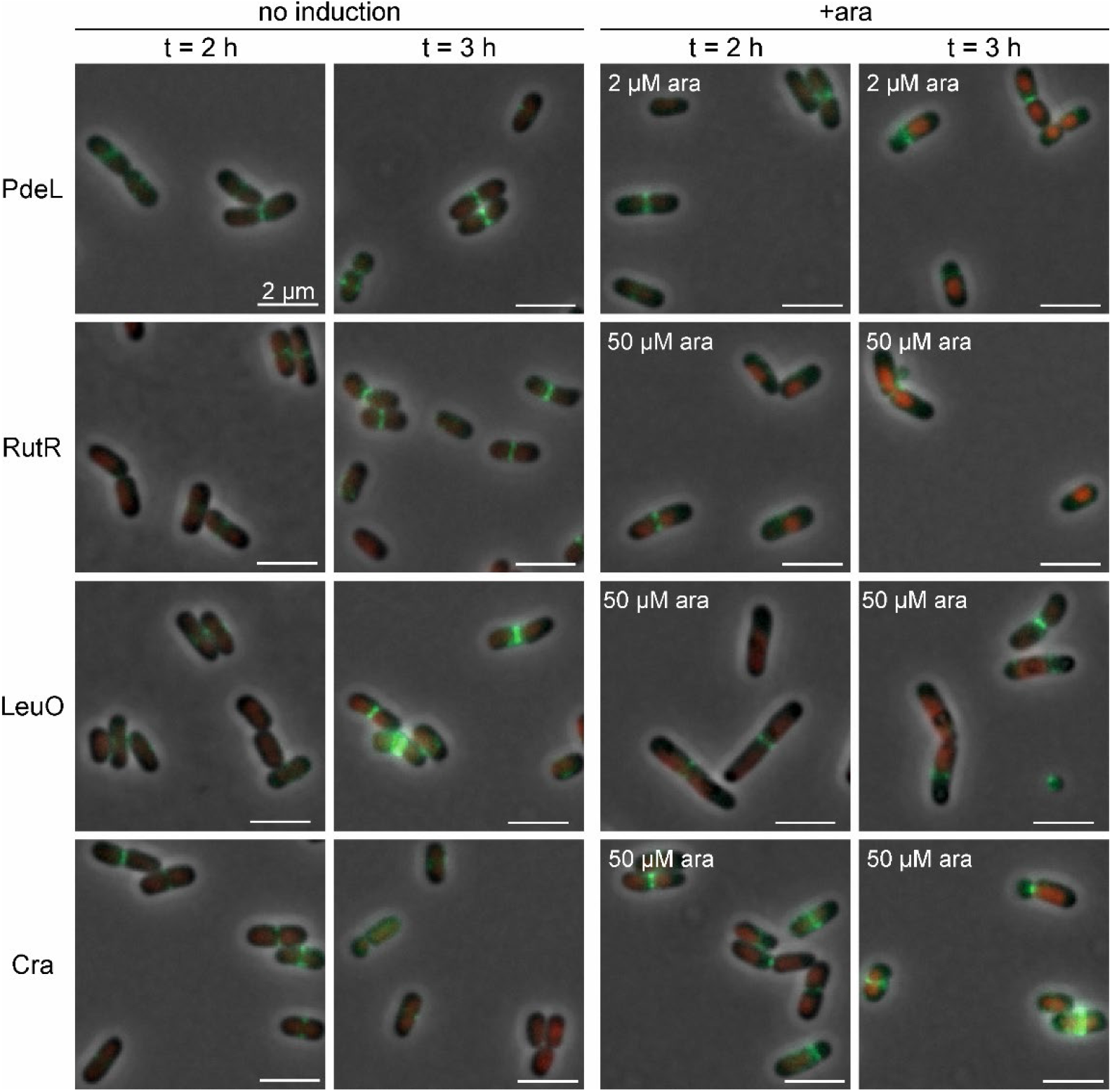
Cell division protein FtsZ is misplaced upon overexpression of transcription regulators RutR, LeuO and Cra. Co-transformants of *hupA-mCherry* Δ*pdeL* strain U159 with plasmids pKECY90 (FtsZ-mNG) and pKECY44 (PdeL), pKECY101 (RutR), pKECY102 (LeuO) and pKECY103 (Cra) were grown in tryptone medium supplemented with chloramphenicol and ampicillin. After one hour of growth *P*_*ara*_ directed expression of transcription regulators genes were induced with L-arabinose, as indicated, and samples were harvested after two (t= 2 h) and three (t = 3h) hours of growth. Representative composite microscopy images with FtsZ-mNG (shown in green) and HupA-mCherry (in red). Full size images are shown in Fig. S10.

Lastly, the amount of PdeL that is synthesized upon induction of *P*_*ara*_ by 2 µM L-arabinose was estimated using purified PdeL-His_6_ as a reference (Fig. S11). After one hour of induction, approximately 40,000 PdeL monomers and after two hours of induction approximately 150,000 monomers of PdeL are present per cell, which corresponds to one PdeL dimer per 250 bp after one hour and four PdeL dimers per 250 bp after two hours of induction (assuming that a single non-replicating nucleoid is present).

## Discussion

Here we have shown that high levels of the transcription regulators PdeL, RcsB, LacI, RutR, Cra, and LeuO lead to nucleoid compaction in *E. coli*. These transcription regulators include single target and local regulators with only one or few specific DNA-binding sites and global regulators with hundreds of DNA-binding sites in the genome (6). Our data suggest that nucleoid compaction is caused by non-specific occupancy of the genomic DNA. Comparable observations of nucleoid compaction have been described recently for the phage T4 protein MotB and for the bacterial DNA-binding toxin SymE (22, 23).

The dual function protein PdeL, a transcription regulator and c-di-GMP specific phosphodiesterase, is nucleoid-associated. Further, ectopic expression of *pdeL* directed by weak induction of *P*_*ara*_ resulted in a high cellular protein level, apparently because PdeL is a protein of high stability. The high level of PdeL caused nucleoid compaction. Likewise, the two-component response regulator RcsB, which carries a FixJ/NarL/LuxR-type DNA-binding domain like PdeL, caused nucleoid compaction, when expressed at similarly high levels as PdeL. Other transcription regulators led to nucleoid compaction as well, and this is independent of the number of specific DNA-binding sites, as shown with the single target regulator LacI, the local regulator RutR, and the global regulators Cra and LeuO (5, 6). In case of LacI nucleoid compaction is independent of specific DNA-binding as shown using the gratuitous inducer IPTG.

The high level of PdeL with approximately 40,000 molecules per cell one hour after induction corresponds to one dimer per 250 bp, while the approximately 150,000 molecules present after two hours of ectopic synthesis would theoretically allow complete coverage of the genome. The cellular levels of the other transcription regulators tested in this study are comparable. Occupancy of the whole genome by a DNA-binding could hinder DNA replication and/or transcription. In case of the toxin SymE it has been shown that toxicity of *symE* overexpression is presumably based on nucleoid condensation and inhibition of DNA as well as RNA synthesis (23). Further, these authors have shown that nucleoid compaction caused by synthesis of SymE at high levels is similar to nucleoid compaction caused by overexpression of H-NS (23, 24). Such a mechanism of genome silencing is possibly utilized by phage T4, which encodes the protein MotB that shortly after infection is synthesized at very high levels corresponding to 40,000 monomers per cell (22). In case of phage T4 MotB, fluorescence imaging using a MotB-GFP fusion demonstrated nucleoid compaction (22) similar to the data shown in this work. Interestingly, MotB has dramatic effects on the transcriptome leading to a relative increase of 1/8 of all transcripts in relation to the whole transcriptome, and of these 1/8 of the transcriptome ∼70% correspond to H-NS repressed genes (22, 25). Thus, phage T4 apparently employs the abundance of MotB to re-program the host transcriptome (22). Remarkably, nucleoid-associated proteins (NAPs) such as H-NS, HU, IHF, and FIS and others are abundant (26), but at their natural level they do not cause nucleoid compaction. H-NS, present at approximately 20,000 molecules per cell binds to the minor groove of AT-rich DNA and forms linear and bridged filaments. HU and IHF are likewise abundant and DNA-bending proteins that presumably contribute to nucleoid structuring and formation of specific regulatory nucleoprotein complexes. FIS is expressed at very high levels with 50,000 molecules per cell during the early exponential growth phase only; it is a DNA-bending protein and organizer of plectonemic structures (7, 27). The cellular levels of the abundant NAPs is apparently well adjusted to their function.

Our data suggest that compaction of the nucleoid by the transcription regulators displaces Z-rings from mid-cell towards the cell poles. Exclusion of the Z-ring from midcell can be an indirect effect mediated by the nucleoid occlusion system and SlmA protein (28). Similarly, the loss of detectable MukB-mNeonGreen foci near the origin of replication can be an indirect consequence of nucleoid compaction, as it is possible that DNA replication is put on hold by excessive occupancy of the genome by overexpressed transcription regulators. In accordance with an indirect effect, the change in positioning of MukB and the FtsZ is observed later than nucleoid compaction.

The serendipitous finding reported here emphasizes the importance of using native protein levels in functional analysis of transcription regulators. Another aspect to be considered when using ectopic expression is that the level of a transcription regulator and the number of its specific DNA-binding sites in the genome is balanced (29). Ectopic expression can be required for a functional analysis, as for example when conditions leading to expression of the gene encoding a transcription regulator and an inducer are not known. In case of transcription regulators that are stable, such as PdeL, ectopic expression even for a short time and with a low inducer concentration may yield far too high protein levels. Thus, if an ectopic expression system is used, the rate and duration of synthesis as well as the protein stability and cellular level should be controlled carefully.

## Material and Methods

### Bacterial strains media and plasmids

E. coli K12 strains and their construction are described in Table 1. Strains were constructed by transduction using phage T4*GT7* or P1 *vir*, and by Lambda-Red-mediated recombineering, respectively (30–33). Plasmids are listed in Table 2 and oligonucleotides used for construction of strains and plasmids are listed in supplementary Table S1. Bacteria were cultured in in LB medium (10 g/l tryptone, 5 g/l yeast extract, and 5 g/l NaCl), tryptone medium (10 g/l tryptone, 5 g/l NaCl), SOB medium (20 g/l tryptone, 5 g/l yeast extract, 0.5 g/l NaCl, 2.5 mM KCl, 10 mM MgCl_2_, pH7.0), or SOC (SOB with 0.4 % glucose). For plates, 15 g/l agar were added. Antibiotics were added to a final concentration of 50 µg/ml ampicillin, 15 µg/ml chloramphenicol, and 15 µg/ml kanamycin, as required; IPTG and L-arabinose, respectively, were added as described in the figures.

### Fluorescence microscopy

For fluorescence microscopy transformants of *E. coli* strain U159 (U65 *hupA-mCherry*_*FRT*_Δ*pdeL*_FRT_) and its derivatives were inoculated to OD_600_ of 0.08 in tryptone medium supplemented with appropriate antibiotics and grown at 37°C whilst shaking. Plasmid *P*_*ara*_ directed expression of the transcription regulators was induced after one hour of growth by adding L-arabinose at the concentration stated in the figures. Bacteria were harvested just before induction (t = 1h), and one and two hours after induction (at t = 2h and t = 3h) by pelleting 500 µl of the culture by centrifugation at 5900 r.c.f. for 1 minute. The bacterial pellets were resuspended in 150 µl of fresh tryptone medium, and 4 µl of these suspensions were spotted onto 1% agarose pads for microscopy. Image acquisition was performed using Zeiss Axio Imager.M2 microscope with an EC Plan-Neofluar 100x/1.30 Oil Ph3 M27 objective. Images were captured and processed using ZEN 2012 software (Carl Zeiss Microscopy GmbH, Germany). Excitation times were 500 ms for PdeL-mVenus fusions, MukB-mNG, FtsZ-mNG and HupA-mCherry. Contrast settings were 1-16384 for phase contrast, 200-2000 for HupA-mCherry, 1-16384 for PdeL-mVenus, 180-300 for MukB-mNG and 180-1000 for FtsZ-mNG, unless otherwise stated in the figures.

### PdeL protein stability analyses

For determining the protein stability of PdeL and its variants, transformants of *E. coli* strain U121 (U65 Δ*pdeL*_FRT_) with plasmids pKECY81 (*P*_*ara*_ *pdeL-3xFLAG*) and pKECY91 (*P*_*ara*_ *pdeL*_*HTH5M*_*-3xFLAG*) were inoculated to an OD_600_ of 0.08 in tryptone medium supplemented with ampicillin. The transformants were grown for 2 hours at 37°C without induction of *P*_*ara*_ to preserve a low protein level for quantification by western blotting. Translation was blocked by adding chloramphenicol to a final concentration of 100 µg/ml. Samples of 2 ml volume were taken just prior and at several time points after inhibition of translation. Bacteria were pelleted by centrifugation and re-suspended in Laemmli buffer for detection of epitope-tagged protein by Western blotting (34)}.

### Protein detection by western blotting and coomassie staining

SDS-PAGE, Coomassie staining, and Western blots were performed as described (34). Unless otherwise described, bacteria equivalent to an OD_600_ of 0.08 were loaded per lane. Equal loading was validated by 2,2,2-trichloroethane (TCE) staining of total protein (35). Epitope-tagged 3xFLAG proteins were detected using primary antibody anti-FLAG M2 from mouse (diluted 1:4,000, catalogue number F3165; Sigma Aldrich, Germany) and Alexa Fluor 680 fluorescent dye-labeled secondary anti-mouse antibody from goat (diluted 1:10,000, catalogue number A21057; Thermo Scientific, Germany). Quantification of protein bands was performed with Odyssey V3.0 software for Western blots and with ImageLab (BioRad, Germany) for Coomassie stained gels.

